# The first complete mitochondrial genome of *Sphaeniscus atilius* (Walker, 1849) (Diptera: Tephritidae) and implications for the phylogenetic relationships of Tephritidae

**DOI:** 10.1101/2023.05.18.541288

**Authors:** Shibao Guo, Junhua Chen, Nan Song, Fangmei Zhang

## Abstract

The nearly complete mitochondrial genome of *Sphaeniscus atilius* was characterized and annotated in this study. The mitogenome was 16,854 bp in length and encoded 37 typical mitochondrial genes, including 13 protein-coding genes, 22 tRNA genes, 2 ribosomal RNA genes, and a control region. The total length of the 13 PCGs was 11,140 bp, and the AT content was 79.8%. There were five types of start codons, ATT (*nad2*, *nad3*, *nad5*, and *nad6*), ATG (*cox2*, *cox3*, *atp6, nad4*, *nad4l*, and *cob*), CGA (*cox1*), as well as ATC (*atp8*) and ATA (*nad1*). Most of the PCGs had typical TAA stop codons, except *nad5* which terminated with incomplete forms T-. Ile, Phe, Leu and Asn were the most frequently used amino acids in mitochondrial PCGs. Most tRNA genes could be folded into the typical cloverleaf structure, except *trnS1* and *trnT* which lacked the dihydrouridine (DHU) and TΨC arms, respectively. Phylogenetic analyses based on 13 protein-coding genes among the available sequenced species of family Tephritidae by maximum likelihood methods suggested the genus relationship of Tephritidae: ((*Bactrocera*, *Dacus*, *Zeugodacus*), *Felderimyia*, *Anastrepha*), (*Acrotaeniostola*, (*Neoceratitis*, *Ceratitis*), *Euleia*, *Rivellia*), (*Procecidochares*, (*Tephritis*, *Sphaenisscus*))))). Our results presented the first mitogenome from *Sphaeniscus* and offer insights into the identification, taxonomy, and phylogeny of *Sphaeniscus atilius*.

## Introduction

Tephritidae, one of the largest families of Diptera, consists of more than 500 genera and almost 5,000 named species and predominantly distributes throughout the temperate and tropical areas of the world [1–3]. This family is also referred to as ‘true’ fruit flies, with hundreds of fruit-eating species accounting for about 40% of the species. It has been reported to attack a great variety of fruit plants, bamboo culms, vegetables, flowers, and seeds [4,5]. In practice, some of the fruit-eating species in the Anastrepha, Bactrocera, Ceratitis, Dacus, and Rhagoletis genera have been considered serious agricultural pests due to their significant economic impact on the production of fruit crops and stored fruit [6,7]. Melon fly, *Zeugodacus cucurbitae* [8] medfly, *Ceratitis fasciventris* [9], together with *Bactrocera latifrons* [10], are well-known examples.

The insect mitochondrial genome has been regarded as a useful molecular marker in studies of phylogenetic and evolutionary analysis, genetic diversity, and species delimitation at the genus or species level, due to its small size, high copy numbers, maternal inheritance, unambiguous orthologous genes, conserved gene composition, and high evolutionary rate [11–14]. Generally, the typical insect mitochondrial genome is a highly conserved circular molecule ranging in size from approximately 14 to 40 kbp, encoding a fixed set of 37 genes, including 13 protein-coding genes (PCGs), 22 transfer RNA (tRNA) genes, two ribosomal RNA (rRNA) genes, and a control region (CR) or the A+T-rich region [14,15], with only a few exceptions. For example, long gene intergenic spacers, gene rearrangements, and gene loss have also been reported in different orders of insects [16–18].

Partial mitochondrial gene sequences have become a preferred approach for inferring phylogenetic and molecular systematic studies in several insect groups, such as *Arma custos* and *Picromerus lewisi* (Hemiptera) [19], *Episymploce splendens* (Blattodea) [18], and *Haematopinus tuberculatus* (Psocodea) [20], *Coomaniella copipes*, *Coomaniella dentata*, and *Dicerca corrugate* (Coleoptera) [21], including the Tephritidae family (*Ceratitis fasciventris* [9], *Bactrocera carambolae* [22], *Bactrocera biguttula* [23], *Zeugodacus cucurbitae* [24], and *Lepidotrigona flavibasis* [25]. The aforementioned studies of mitochondrial gene sequences explored the origin and evolution of insects, explained the species and evolution of the system, and revealed the geographical distribution of intraspecific polymorphism.

*Sphaeniscus atilius* (Diptera: Tephritidae: *Sphaeniscus*), which is distributed from India to Russia, Korea, Japan, Australasian and Oceanian regions, can be morphologically distinguished from any other tephritid species based on clear diagnostic morphological features, including an almost entirely dark brown body, the number of orbital frontal setae, and an almost perpendicular base of the discal band of the wing [26]. To date, the complete mitochondrial genomes (mitogenomes) of 46 species, belonging to 14 genera of Tephritidae, are available in GenBank (https://www.ncbi.nlm.nih.gov/nucleotide/). All of these species belong to 5 subfamilies, Sciomyzidae, Sepsidae, Lauxaniidae, Celyphidae, Platystomatidae, and Tephritidae, respectively. However, there are no reports on the molecular phylogeny studies of the mitochondrial genome information in *S. atilius*, which limits our comprehensive understanding of the evolutionary and phylogenetic relationships of *S. atilius*.

In the current study, we sequenced, annotated, and described the complete mitogenome of *S. atilius* using next-generation sequencing, which is the first complete mitogenome sequence reported in the genus *Sphaeniscus*. We predicted and analyzed the gene organization, base composition, protein-coding genes (PCGs), codon usage, and the structure of the transfer RNAs (tRNAs) and ribosomal RNAs (rRNAs) of its mitochondrial genome. Additionally, we carried out phylogenetic analyses based on Maximum Likelihood (ML) to assess the phylogenetic position of *S. atilius*. These results will be greatly helpful for clarifying the phylogenetic status and relationships between different species of Tephritidae.

## Materials and Methods

### Taxon sampling and DNA extraction

Specimens of *S. atilius* were collected in Mount Jigong, Xinyang, Henan Province, China (31°48′43″N, 114°05′43″E), in June 2020. Specimens were preserved in 100% ethanol, and stored at -20°C. After morphological identification, total genomic DNA was extracted from muscle tissue of pre-thoraxes using Tissue DNA kit (TIANGEN Biotech, Beijing, China) according to the manufacture’s protocol. The impurities and concentration were detected by agarose (1%) electrophoresis and Nanodrop spectrophotometer (ThermoFisher Scientific, Waltham, MA), respectively.

### Sequencing and Assembling of Mitochondrial Genome

TruSeq library was prepared an insert size of 400 bp, and this was sequenced on an Illumina HiSeq 2500 platform (150 bp paired-end reads). Quantity of sequencing data for each sample was at least 20 Gb. The sequencing data was detected and filtered by software NGS QC-Toolkit v2.5 [27]. The sequencing was performed at Beijing Novogene Bioinformatics Technology Co., Ltd, China. As a result, about 2 Gb raw paired reads were generated. After removing the connector and the unmatched, short, and poor-quality reads, the high-quality reads (Q20 > 90% and Q30 > 80%) were used for genome assembly. De novo assembly was performed with IDBA-UD v. 1.1.1 [28]. The parameter settings for assembly were the following: 200 for the setting of minimum size of contig, and an initial k-mer size of 41, an iteration size of 10, and a maximum k-mer size of 91.

### Mitochondrial Genome Annotation and Analysis

Preliminary annotation of mitochondrial genome was conducted using MITOS (http://mitos.bioinf.uni-leipzig.de/index.py) [29]. The gene boundaries were refined by blasting against closely related species. The secondary structures were predicted in MITOS and redrawn in Adobe Illustrator. The mitogenome maps were drawn using OGDRAW v1.3.1 (http://ogdraw.mpimp-golm.mpg.de/cgi-bin/ogdraw.pl) [30]. The annotated mitogenome sequence of *S. atilius* has been submitted to GenBank with accession number OQ909100.

The base composition and relative synonymous codon usage (RSCU) were analyzed and computed using MEGA 7.0 [31]. AT and GC skews were calculated following the formula: AT Skew=(A − T)/(A + T), GC Skew=(G − C)/(G + C) [32]. The nucleotide diversity (Pi) and nonsynonymous (Ka)/synonymous (Ks) mutation rate ratios were calculated by DnaSP v5.10.01 [33].

### Phylogenetic Analysis

The new mitogenome sequence of *S. atilius* was merged with the existing dipteran mitogenome sequences. The amino acid sequences of 13 PCGs 43 species from three superfamilies belonging to 2 families and 13 genus of Tephritidae (Table 1) were aligned individually using MAFFT with default parameters [34]. The alignments were trimmed by trimAl [35]. Then, the resulting alignments were concatenated using SequenceMatrix 1.8 [36]. Phylogenetic analysis was conducted using Maximum Likelihood (ML) inference method. ML tree was searched in IQ-TREE v2.2.3 [37]. The best-fitting model for the alignment was estimated by ModelFinder [38]. Branch support was assessed by 10,000 ultrafast bootstrap replicates [39].

**Table 1.**
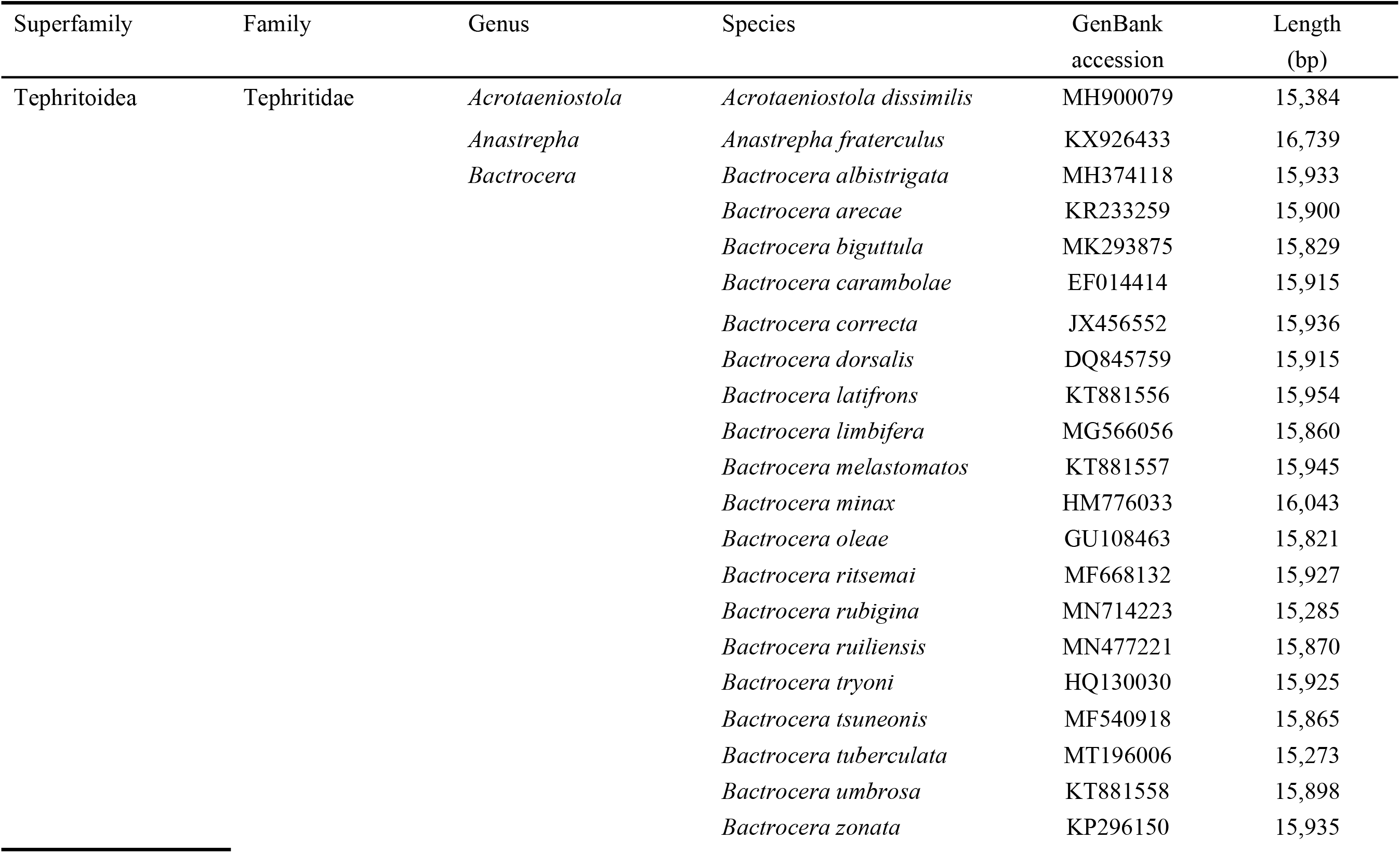
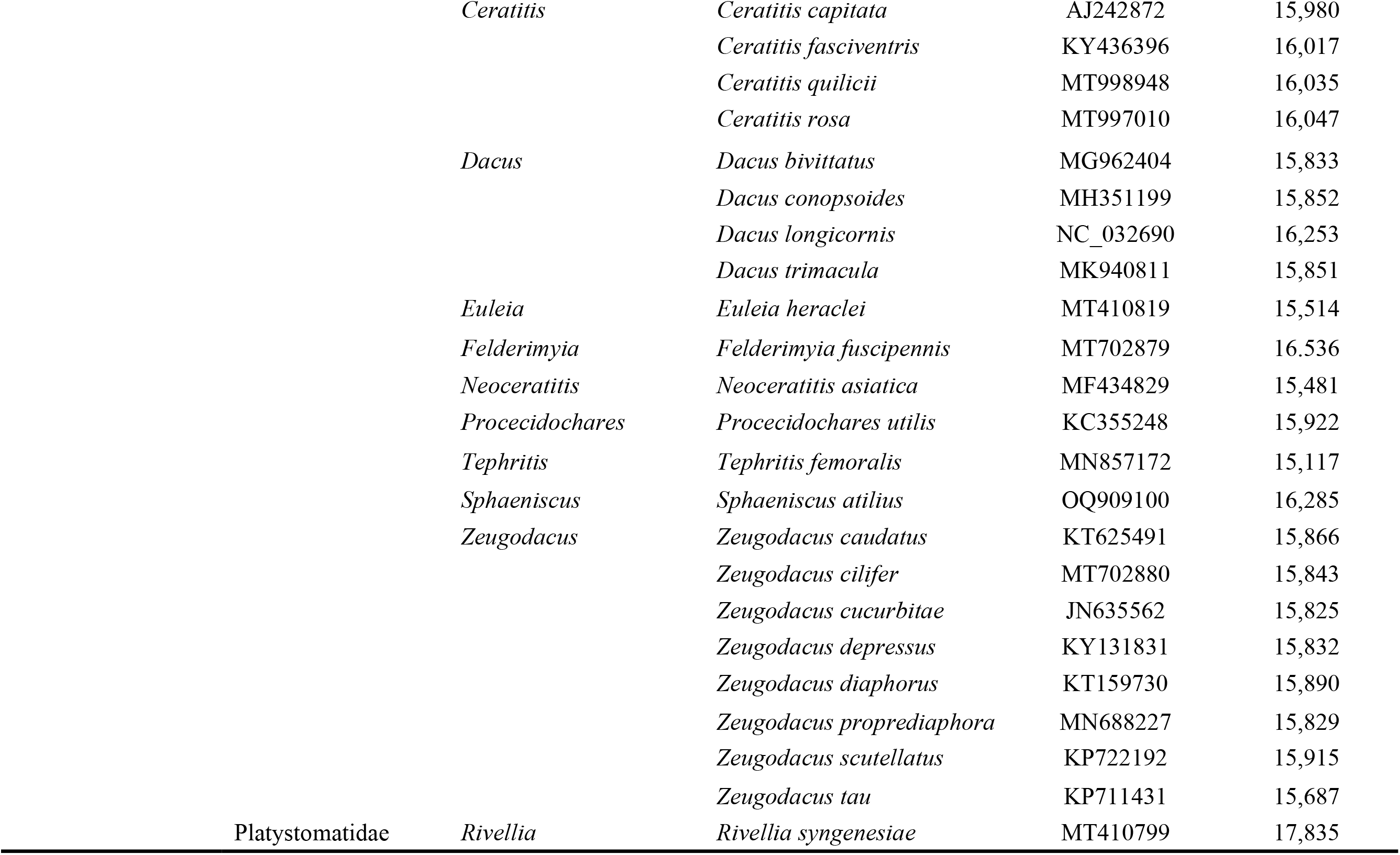
List of taxa used for phylogenetic.

## Results

### Genome organization

The length of mitogenome in *S. atilius* was 16,854 bp in length with typical circular molecules, which consisted of 37 genes, including 13 PCGs, 22 tRNA genes, and two rRNA genes (Figure 1, Table 2). In addition, a major non-coding region (D-loop) known as the control region (CR) or A+T-rich region was found between *rrnS* and *trnI*. The heavy chain (H-strand) encoded 23 genes (9 PCGs and 14 tRNAs). The other (4 PCGs, 8 tRNAs and 2 rRNAs) were transcribed from the light chain (L-strand).

**Figure 1.**
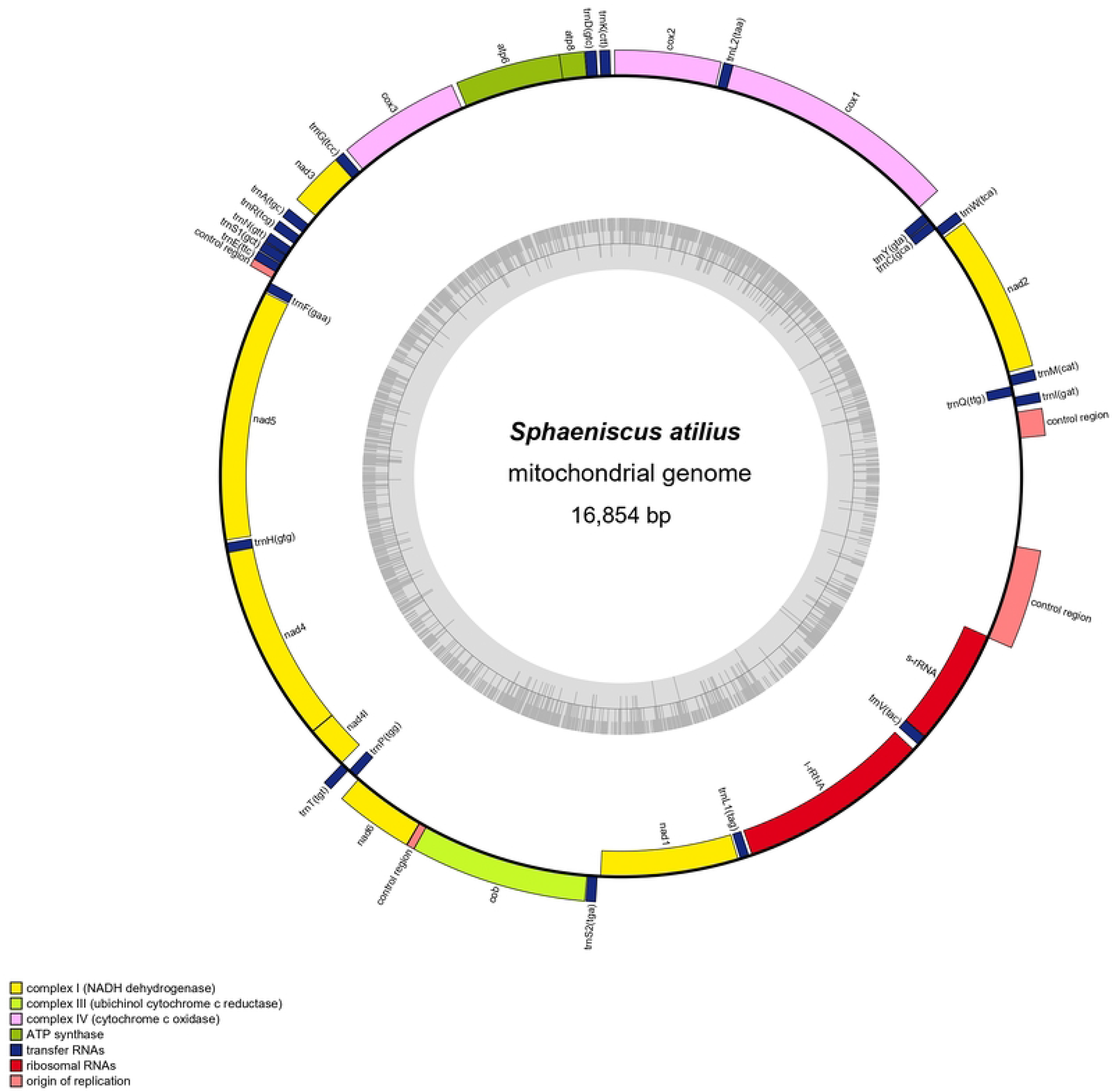
The mitochondrial genome of *S. atilius* showing the protein coding genes, tRNAs, rRNAs and non-coding regions

**Table 2.** Summary of the mitogenome of *S. atilius*.

| Gene | Strand | Location | Size (bp) | Anti codon | Start codon | Stop codon | Intergenic nucleotides |
| --- | --- | --- | --- | --- | --- | --- | --- |
| *CR1* |  | 255-432 | 178 |  |  |  | 0 |
| *trnI* | H | 469-535 | 67 | GAT |  |  | 7 |
| *trnQ* | L | 543-611 | 69 | TTG |  |  | -1 |
| *trnM* | H | 611-682 | 72 | CAT |  |  | 21 |
| *nad2* | H | 704-1711 | 1008 |  | ATT | TAA | 14 |
| *trnW* | H | 1726-1793 | 68 | TCA |  |  | -8 |
| *trnC* | L | 1786-1851 | 66 | GCA |  |  | 1 |
| *trnY* | L | 1853-1919 | 67 | GTA |  |  | 47 |
| *cox1* | H | 1967-3502 | 1536 |  | CGA | TAA | -5 |
| *trnL2* | H | 3498-3463 | 66 | TAA |  |  | 10 |
| *cox2* | H | 3574-4257 | 684 |  | ATG | TAA | 23 |
| *trnK* | H | 4281-4351 | 71 | CTT |  |  | 16 |
| *trnD* | H | 4368-4441 | 74 | GTC |  |  | 0 |
| *atp8* | H | 4442-4606 | 165 |  | ATC | TAA | -7 |
| *atp6* | H | 4600-5277 | 678 |  | ATG | TAA | 28 |
| *cox3* | H | 5306-6091 | 786 |  | ATG | TAA | 22 |
| *trnG* | H | 6114-6179 | 66 | TCC |  |  | 0 |
| *nad3* | H | 6180-6533 | 354 |  | ATT | TAA | 72 |
| *trnA* | H | 6606-5574 | 69 | TGC |  |  | 27 |
| *trnR* | H | 6702-6769 | 68 | TCG |  |  | 28 |
| *trnN* | H | 6798-6862 | 65 | GTT |  |  | 0 |
| *trnS1* | H | 6863-6929 | 67 | GCT |  |  | 2 |
| *trnE* | H | 6932-6997 | 66 | TTC |  |  | -8 |
| *CR2* |  | 6990-7043 | 54 |  |  |  | 0 |
| *trnF* | L | 7072-7139 | 68 | GAA |  |  | 6 |
| *nad5* | L | 7146-8859 | 1714 |  | ATT | T(AA) | 15 |
| *trnH* | L | 8875-8940 | 66 | GTG |  |  | 0 |
| *nad4* | L | 8941-10281 | 1341 |  | ATG | TAA | -1 |
| *nad4L* | L | 10281-10571 | 291 |  | ATG | TAA | 2 |
| *trnT* | H | 10574-10637 | 64 | TGT |  |  | 0 |
| *trnP* | L | 10638-10703 | 66 | TGG |  |  | 20 |
| *nad6* | H | 10724-11230 | 507 |  | ATT | TAA | -2 |
| *CR3* |  | 11229-11284 | 56 |  |  |  | 0 |
| *cob* | H | 11279-12415 | 1137 |  | ATG | TAA | 2 |
| *trnS2* | H | 12418-12484 | 67 | TGA |  |  | 17 |
| *nad1* | L | 12502-13440 | 939 |  | ATA | TAA | 10 |
| *trnL1* | L | 13451-13516 | 66 | TAG |  |  | 16 |
| *16sRNA* | L | 13533-14841 | 1309 |  |  |  | 41 |
| *trnV* | L | 14883-14954 | 72 | TAC |  |  | -1 |
| *12sRNA* | L | 14954-15752 | 799 |  |  |  | -12 |
| *CR4* |  | 15741-16385 | 645 |  |  |  | 0 |

Strand of the genes is presented as L for majority and H for minority strand. In the column for intergenic length, a positive sign indicates the interval in base pairs between genes, while the negative sign indicates overlapping base pairs between genes.

There were five intergenic overlapping regions totaling 29 bp, with varying lengths of 1-8 bp, which mainly occured in tRNA genes. The two longest overlapping regions, both with a length of 8 bp, occur between *nad2* and *trnW* and between *trnS1* and *trnE*. Sixteen intergenic spacer regions were examined, totaling 339 bp in length, with the longest spacer sequence (72 bp) located between *trnG* and *nad3*, followed by a 47 bp spacer between *trnC* and *trnY*. There were also three regions without gene overlaps or intergenic spacers.

The base nucleotide composition of the mitogenome is 41.80% A, 39.92% T, 7.67% G, and 10.61% C, respectively. The content of A+T (81.72%) was higher than that of G+C (18.28%). The full mitogenome of *S. atilius* exhibited a positive AT skew (0.02) and a negative GC skew (-0.16) (Table 3).

**Table 3.**
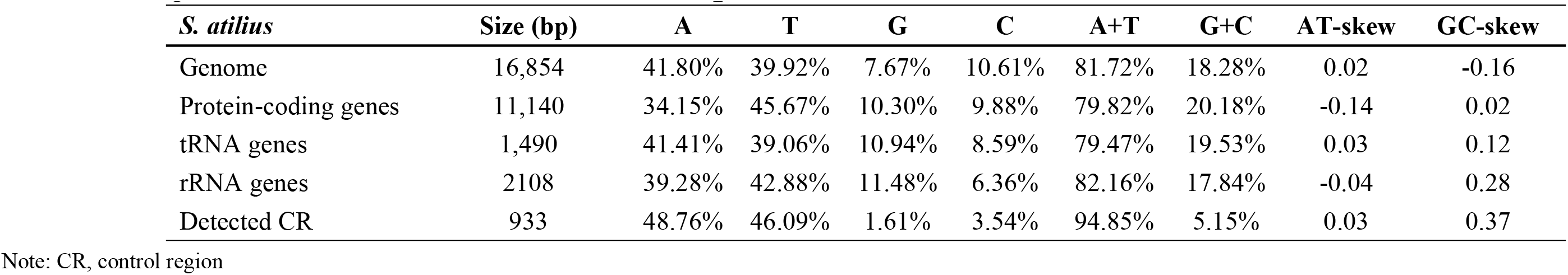
Composition and skewness of the *S. atilius* mitogenome.

### Protein-coding Genes

The total length of the 13 PGGs of in the *S. a*tilius mitogenome was 11,140 bp, accounting for 66.10% of the whole mitogenome sequence (16,854 bp), and encoding a total of 3,612 codons. Among them, *nad5* (1,714 bp) was found to be the longest sequence, and *nad4L* (291 bp) was the shortest (Table 2). Nine PCGs (*nad2*, *cox1*, *cox2*, *atp8*, *atp6*, *cox3*, *nad3*, *nad6*, and *cob*) were coded on the H-strand, while the remaining four PCGs (*nad5*, *nad4*, *nad4L*, and *nad1*) were located on the L-strand. The content of AT and GC was found to be 79.82% and 20.18% in the 13 PCGs, exhibiting a highly AT bias (Table 3). The AT skew was negative (-0.14), while the GC skew was positive (0.02) in the PCGs. Additionally, six PCGs (*cox2*, *cox3*, *atp6*, *nad4*, *nad4L*, and *cob*) initiated with an ATG start codons, four PCGs (*nad2*, *nad3*, *nad5* and *nad6*) used ATT as the start codon, nad1 used ATA, *atp8* used ATC, and *cox1* used CGA. The termination codons of 12 PCGs were TAA, and the remaining genes (*nad5*) used an incomplete stop codon T.

The amino acid usage of the 13PCGs and the relative synonymous codon usage (RSCU) frequency statistical analysis results are shown in Figures 2 and Table 4. In total, the mitogenome contained 3,885 codons, excluding stop codons (172 bp). The most frequently used amino acids in mitochondrial PCGs were Ile, Phe, Leu, and Asn, accounting for 16.68%, 12.51%, 10.86%, and 8.13% of the total amino acids, respectively (Figure 2A). The most frequently used codons were UUU (436), AUU (436), UUA (342), and AAU (277). The relatively scarce amino acids were Met (0.62%), Trp (0.67%), Arg1 (0.93%), and Asp (1.23%), and the frequency of the codons CUC, CUG, and CGC was zero. The most frequent synonymous codons were UUA, AGA, and GUU, with UUA having the highest frequency of relative synonymous codons (RSCU = 4.86). Conversely, CUC, CUG, and CGC had a relatively low frequency of synonymous codons, with an RSCU of 0 (Figure 2B).

**Figure 2.**
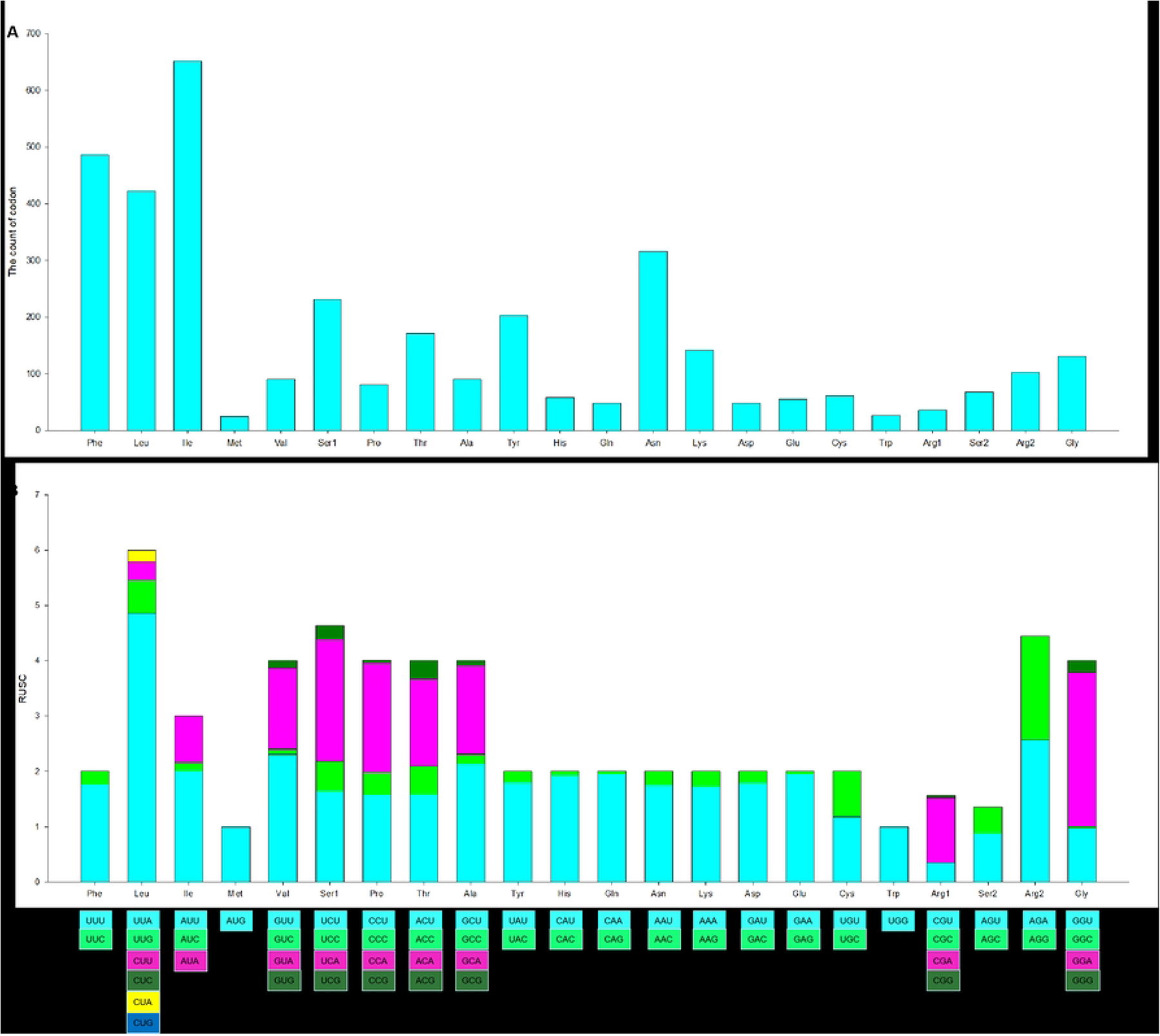
Usage of amino acids of 13 PCGs in the *S. atilius* mitogenome. (A) The total numbers of codon families; (B) Relative synonymous codon usage (RSCU) of codon families

**Table 4.** Codon number and RSCU in *S. atilius* mitochondrial PCGs.

| Codon | Count | RSCU | Codon | Count | RSCU | Codon | Count | RSCU | Codon | Count | RSCU |
| --- | --- | --- | --- | --- | --- | --- | --- | --- | --- | --- | --- |
| UUU(F) | 431 | 1.77 | UCU(S) | 82 | 1.65 | UAU(Y) | 183 | 1.8 | UGU(C) | 36 | 1.18 |
| UUC(F) | 55 | 0.23 | UCC(S) | 27 | 0.54 | UAC(Y) | 20 | 0.2 | UGC(C) | 25 | 0.82 |
| UUA(L) | 342 | 4.86 | UCA(S) | 109 | 2.19 | UAA(*) | 60 | 1.05 | UGA(*) | 93 | 1.62 |
| UUG(L) | 42 | 0.6 | UCG(S) | 13 | 0.26 | UAG(*) | 19 | 0.33 | UGG(W) | 26 | 1 |
| CUU(L) | 23 | 0.33 | CCU(P) | 32 | 1.58 | CAU(H) | 56 | 1.93 | CGU(R) | 8 | 0.35 |
| CUC(L) | 0 | 0 | CCC(P) | 8 | 0.4 | CAC(H) | 2 | 0.07 | CGC(R) | 0 | 0 |
| CUA(L) | 15 | 0.21 | CCA(P) | 40 | 1.98 | CAA(Q) | 47 | 1.96 | CGA(R) | 27 | 1.17 |
| CUG(L) | 0 | 0 | CCG(P) | 1 | 0.05 | CAG(Q) | 1 | 0.04 | CGG(R) | 1 | 0.04 |
| AUU(I) | 436 | 2.01 | ACU(T) | 68 | 1.59 | AAU(N) | 277 | 1.75 | AGU(S) | 44 | 0.88 |
| AUC(I) | 33 | 0.15 | ACC(T) | 22 | 0.51 | AAC(N) | 39 | 0.25 | AGC(S) | 24 | 0.48 |
| AUA(I) | 183 | 0.84 | ACA(T) | 67 | 1.57 | AAA(K) | 123 | 1.73 | AGA(R) | 59 | 2.57 |
| AUG(M) | 24 | 1 | ACG(T) | 14 | 0.33 | AAG(K) | 19 | 0.27 | AGG(R) | 43 | 1.87 |
| GUU(V) | 52 | 2.31 | GCU(A) | 48 | 2.13 | GAU(D) | 43 | 1.79 | GGU(G) | 32 | 0.98 |
| GUC(V) | 2 | 0.09 | GCC(A) | 4 | 0.18 | GAC(D) | 5 | 0.21 | GGC(G) | 1 | 0.03 |
| GUA(V) | 33 | 1.47 | GCA(A) | 36 | 1.6 | GAA(E) | 54 | 1.96 | GGA(G) | 91 | 2.78 |
| GUG(V) | 3 | 0.13 | GCG(A) | 2 | 0.09 | GAG(E) | 1 | 0.04 | GGG(G) | 7 | 0.21 |

In addition, the nucleotide diversity (Pi) of the PCGs among 46 species was calculated, ranging from 0.14 to 0.22 (Figure 3). Among them, *nad6* (Pi=0.22) showed the most diverse nucleotide variability among all PCGs, followed by *nad2* (Pi=0.21), *nad3* (Pi=0.19), and *atp8* (Pi=0.18). The *nad5* (Pi=0.14), *nad1* (Pi=0.15), and *cox3* (0.15) genes exhibited relatively low values of nucleotide variability. The ratio of Ka/Ks was calculated for each gene of the 13 PCGs (Figure 3). The value of the *cox3* gene (Ka/Ks=1.26) was higher than others. Meanwhile, the ratio of Ka/Ks of the other 12 PCGs were all significantly less than 1, with the value of the *cox1* gene being the lowest (Ka/Ks=0.06).

**Figure 3.**
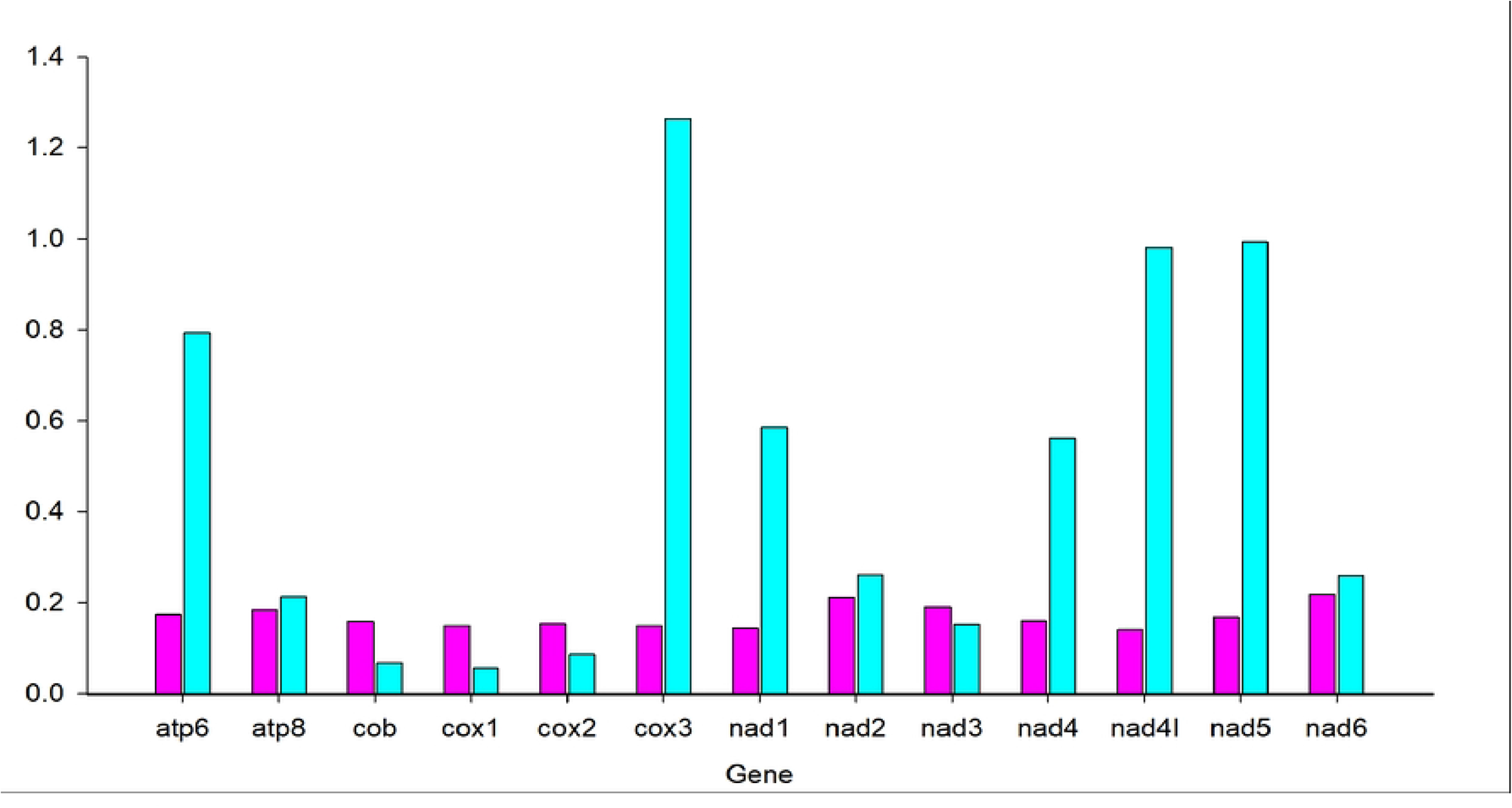
The nucleotide diversity (Pi) and nonsynonymous (Ka) /synonymous (Ks) substitution rate ratios of 13 PCGs of Tephritidae species

### tRNAs and rRNAs

The 22 tRNA genes of the mitogenome of *S. atilius* had 1,490 bp in length, 9.51% of the entire mitogenome, ranging from 65 bp (*trnN*) to 74 bp (*trnD*) (Table 2). Among these, 14 genes (*trn1, trnM, trnW, trnL2, trnK, trnD, trnG, trnA, trnR, trnN, trnS1, trnE, trnT*, and *trnS2*) were located on the H-strand and other 8 genes (*trnQ*, *trnC*, *trnY*, *trnF*, *trnH*, *trnP*, *trnL1*, and *trnV*) were located on the L-strand. Through the analysis of the secondary structure of the tRNAs (Figure 4), most tRNA genes could be folded into the typical cloverleaf secondary structure, while *trnS1* and *trnT* lacked the dihydrouridine (DHU) and TΨC arms, respectively. In the secondary structures of all tRNAs of *S. atilius* (Fig. 3), 3 or 4 base paris in the DHU arms, and 4 or 5 base paris in the TΨC arms. Except the classic base pairs (A-U and C-G), fourteen wobble base pairs (G-U) were detected in 9 genes (*trnA*, *trnC*, *trnF*, *trnG*, *trnH*, *trnP*, *trnQ*, *trnT*, and *trnV*), which occur in the amino acid-accepting arms, anticodon arms, TψC arms, or DHU arms. Of them, *trnH* had the highest rate (three pairs each). Besides, five pairs of U-U base mismatches (in *trnA*, *trnG*, *trnL1*, *trnR*, *trnV*, and *trnW*) and 1 other mismatches base (in *trnN*) were found in the TψC arms. The AT and GC content are 79.47% and 19.53% in the 22 tRNA genes, respectively, with a positive AT skew (0.03) and GC skew (0.12) (Table 3).

**Figure 4.**
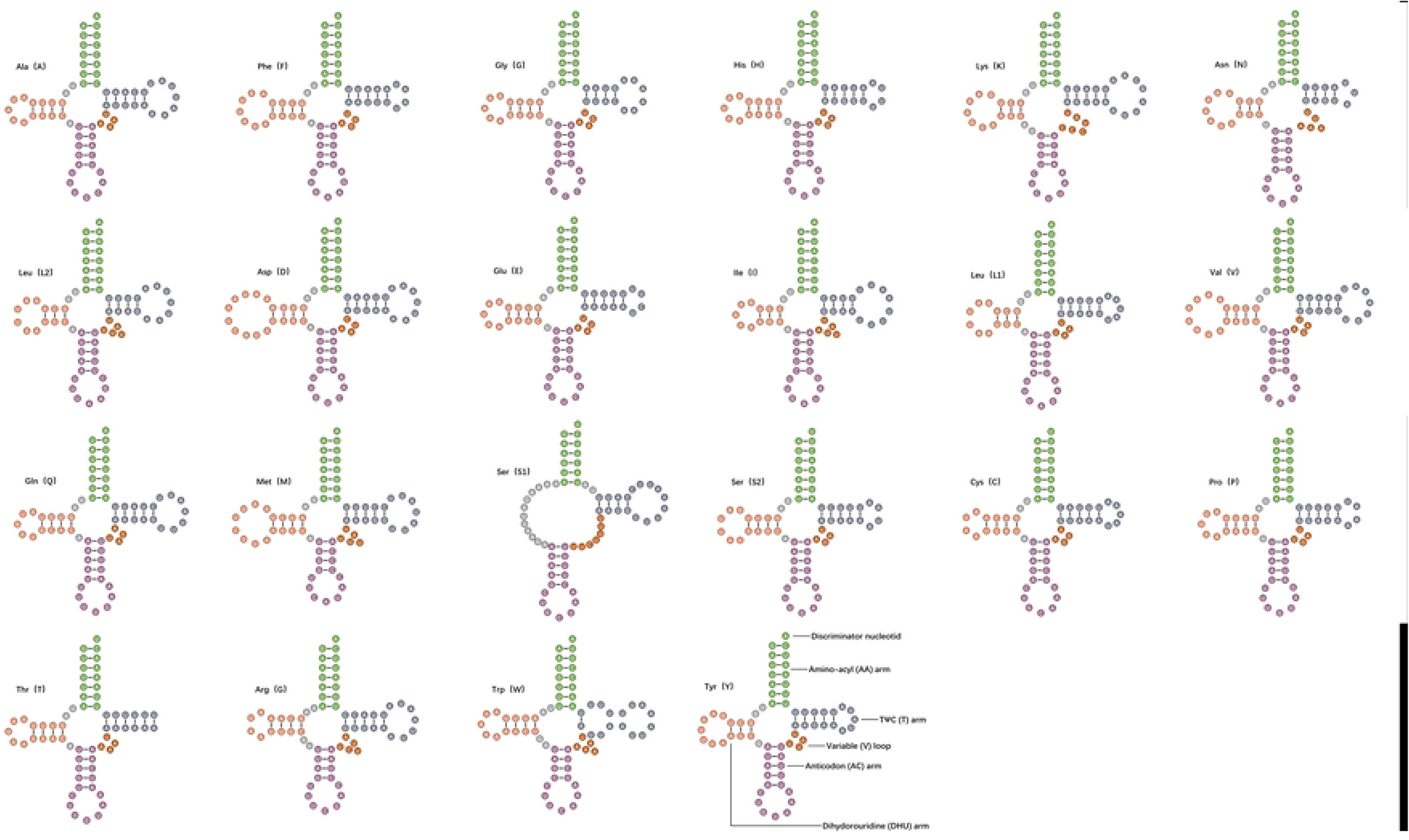
Secondary structures of the transfer RNA genes (tRNAs) in *S. atilius* mitogenome

There were two rRNAs in the mitogenome of *S. atilius*: a 1,309-bp *16S rRNA* (*rrnL*) and a 799-bp *12S rRNA* (*rrnS*) (Table 2). The *16S rRNA* subunit was located between *trnL1* and *trnV*, while the *12S rRNA* gene was located between *trnV* and the control region, embedded in the L-strand. The AT content was 82.16%, with a negative AT skew (-0.04) and positive GC skew (0.28) (Table 3).

### Control Region

The mitogenome of *S. atilius* contained four large non-coding regions, CR1 (178 bp), CR2 (54 bp), CR3 (56 bp) and CR4 (645 bp), and located between the *rrnS* and *trnI*, *trnE* and *trnF,* nad6 and *cob*, and *rrnS* and *trnI*. The AT content of these non-coding regions (94.85%) was higher than that of the whole mitogenome (81.72%), PCGs (79.82%), tRNAs (79.47%) and rRNAs (82.16%). Additionally, both positive AT skew (0.03) and GC skew (0.37) were detected in the control region (Table 3).

### Phylogenetic Analysis

Phylogenetic tree was constructed based on the nucleotide sequences of the 13 PCGs from the mitogenomes of *S. atilius* and other 43 species of Tephritidae using ML method (Figure 5). The result revealed that the thirteen genera of Tephritidae species followed the following monophyletic relationships: ((*Bactrocera*, *Dacus*, *Zeugodacus*), *Felderimyia*, *Anastrepha*), (*Acrotaeniostola*, (*Neoceratitis*, *Ceratitis*), *Euleia*, *Rivellia*), (*Procecidochares*, (*Tephritis*, *Sphaenisscus*))))). Of them, *Bactrocera* (19 exemplars) formed a separate clade at the top of phylogenetic tree, and formed the sister group of a clade including *Dacus* (4 exemplars), *Zeugodacus* (8 exemplars), *Felderimyia* (1 exemplar) and *Anastrepha* (1 exemplar). Ceratitis (4 exemplars) formed a monophyletic group and was sister to *Acrotaeniostola* (1 exemplar) and *Neoceratitis* (1 exemplar). *Euleia* (1 exemplar) and *Rivellia* (1 exemplar) formed the separate clades, respectively. *Sphaenisscus* (1 exemplar) and *Tephritis* (1 exemplar) clustered together, and formed a sister group to *Procecidochares* (1 exemplar). *S. atilius* and *T. femoralis* were closely related and formed a sister group to *Procecidochares utilis*.

**Figure 5.**
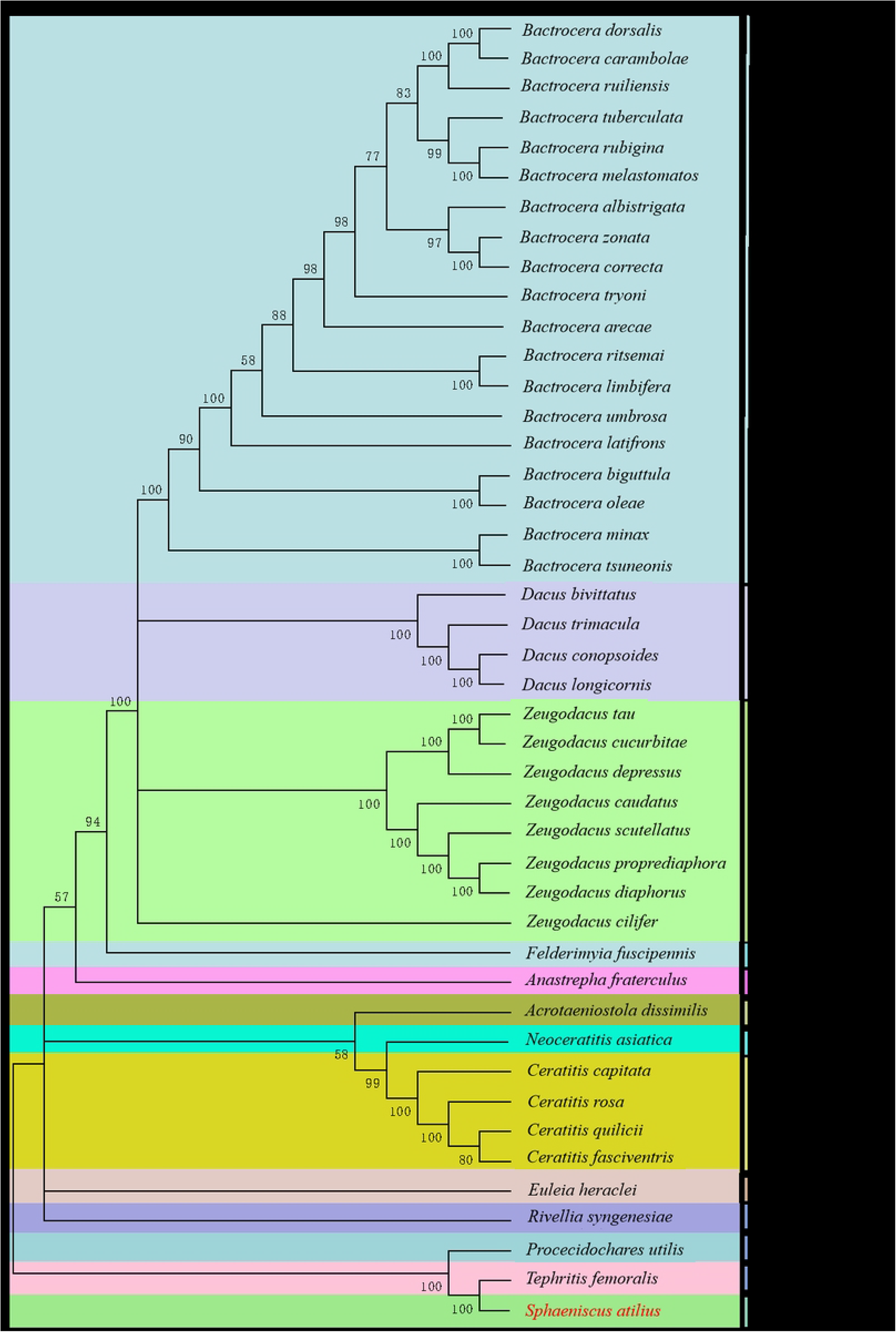
Phylogenetic reconstruction of 43 PCGs sequences was performed using the maximum likelihood analysis. The species sequenced in this study are indicated with red.

## Disscusion

The complete mitogenome of *S. atilius* was a circular, double-stranded DNA molecule with a total length of 16, 854 bp, which was similar to that of other Tephritoidea insects analyzed, ranging from 15,117 bp (*Tephritis femoralis*) to 16,739 bp (*Anastrepha fraterculus*) (Table 1). It has the typical organization and composition of an insect mitochondrion, containing 37 genes, including 13 protein-coding genes (PCGs), 22 tRNA genes, and 2 rRNA genes, which was consistent with those described in other subfamily Tephritoidea insects, such as *Bactrocera carambolae* [22], *Zeugodacus cucurbitae* [24], *Dacus haikouensis* [40]. The nucleotide composition of all regions showed a strong AT bias [41], as seen in other insects [18, 19, 42]. The AT-skew of the full mitogenome is positive (0.02), while the GC skew is negative (-0.16), indicating that the content of bases C is higher than that of G, and A is higher than T in the whole genome.

The mitogenome commonly exhibited compact arrangements, such as small intergenic spacers, or overlapping genes [43]. In our present study, intergenic overlapping regions ranging from 1 to 8 bp, with a total length of 29 bp, were examined. Overlapping regions of similar size are common among Tephritoidea insects [9, 10, 22, 23], while their positions vary. For example, the longest overlaps were found between *atp8* and *atp6* in *B. carambolae* [22], and between *nad2* and *trnW*, and between *trnS1* and *trnE* in *S. atilius*. Most of the gene overlaps occurred in tRNA genes, possibly due to the lower evolutionary constraints of these genes [44].

Sixteen intergenic spacer regions were examined, with a total length of 339 bp, and the longest spacer sequence (72 bp) was located between *trnG* and *nad3*. Larger intergenic spacers in mitogenomes had been reported in Tephritoidea insects, such as, 94 bp in *Bactrocera latifrons*, 82 bp in *Bactrocera melastomatos* and 79 bp in *Bactrocera umbrosa* [10]. Generally, the duplication/random loss model and slipped-strand mispairing have been used to explain the origin of mitogenome intergenic spacers [16, 45]. Whether these long spacer regions were functional was controversial [18].

The 13 PCGs of the *S. atilius* mitogenome were found to be 11,140 bp in length and used a variety of start codons, including ATG for *cox2*, *cox3*, *atp6*, *nad4*, *nad4L*, and *cob*; ATT for *nad2*, *nad3*, *nad5*, and *nad6*; ATA for *nad1*; CGA for *cox1*; and ATC for *atp8*, as reported for *Bactrocera biguttula* [23] and *Bactrocera carambolae* PCGs [22]. Alternative start codons have also been found in other insects, such as TTG in *Arma custos* and *Picromerus lewisi* [19] and GTG in *Nisia fuliginosa* [42]. The *cox1* gene in the *S. atilius* mitogenome used CGA as the starting codon, consistent with other known insects [46]. However, the starting codon of *cox1* was not always uniform, for example, TTG in *Episymploce splendens* [18] and ATA in *Anastatus fulloi* [47]. The typical termination codons TAA was employed in 12 PCGs, which is common among metazoans [18, 47], with one exception of an incomplete stop codon T for *nad5*. This exception was commonly observed in arthropod mitogenomes [21, 47], and may be attributed to post-transcriptional modification during the mRNA maturation process [43, 48].

The four most frequently used codons, UUU (Phe), AUU (Ile), UUA (Leu), and AAU (Asn) were observed in the *S. atilius* mitogenome, which was similar to other insect mitogenomes, such as those of Ephemeroptera [49], Coleoptera [50] and Hemiptera [14]. Meanwhile, the RSCU analysis of the PCGs also indicated that A and U were the components that contributed to the high A + T bias of the full mitogenome. The nucleotide diversity (Pi) and the ratio of *Ka/Ks* of the PCGs among 43 Tephritoidea species were calculated. The results showed that *nad6* exhibited the most diverse nucleotide variability among all PCGs, while *nad5*, *nad1*, and *cox3* exhibited a relatively low variation rate and were the most conserved genes. The overall ratios of *Ka/Ks* for most PCG genes were significantly less than 1, which suggests that these PCGs were under purifying selection. The *cox1* gene had the lowest *Ka/Ks* ratio (*Ka/Ks*=0.06), indicating that this gene had a relatively slow evolutionary rate [51]. This phenomenon occurred in almost all animals [52], and has been subjected to species identification and evolutionary analysis in various arthropod species in Tephritoidea, as well as like in other insects [24, 53].

As in other insects, most tRNA genes in *S. atilius* could be folded into the typical clover-leaf secondary structure, while *trnS1* lacked the dihydrouridine (DHU) arm and *trnT* lacked the TΨC arm. This feature of *trnS1* has been observed in many other insect mitogenomes [21, 42, 44, 47, 54]. In addition, 14 wobble base pairs (G-U) and 5 pairs of U-U base mismatches in the tRNA genes of the *S. atilius* mitogenome were observed. Previous reports have suggested that wobble and mismatched pairs, which commonly occur in insect tRNAs, were usually corrected through the editing process and sustain the transport function [55, 56]. The length, location, and base composition of the two rRNA genes were similar to those of other Tephritoidea insects, such as *Bactrocera arecae* [57] and *Bactrocera biguttula* [23].

In addition, the *S. atilius* mitogenome contained four putative control regions, known as the A+T-rich region, which was supposed to act as the origin of genome replication and gene transcription [15, 48] and ranges in size from tens to several thousands of base pairs [58–60]. The control region was a source of length variation in the mitogenomeis [58, 61]. In the *S. atilius* mitogenome, 933 bp in length were sequenced, which was far less than the extremely long control region of 4,651 bp in *Picromerus lewisi* [19], and the longest known control region length of 1,141 bp in Tephritidae insects [62]. The control region was located between 12sRNA (*rrnS*) and *trnI* genes and has a higher AT content (94.85%), compared to that of Tephritidae insects, such as *Bactrocera arecae* (86.0%) [57], *Bactrocera melastomatos* (89.0%) [10], and *Ceratitis fasciventris* (90.24%) [9].

In the present study, the phylogenetic relationships within Tephritidae can be presented as follows: ((*Bactrocera*, *Dacus*, *Zeugodacus*), *Felderimyia*, *Anastrepha*), (*Acrotaeniostola*, (*Neoceratitis*, *Ceratitis*), *Euleia*, *Rivellia*), (*Procecidochares*, (*Tephritis*, *Sphaenisscus*))))), in concordance with the findings of previous phylogenetic studies [23, 24, 63, 64]. The results showed that four genera (*Felderimyia*, *Zeugodacus*, *Dacus* and *Bactrocera*) are closely related and they gathered to form a sister group. *Dacus* and *Zeugodacus* constituted a sister clade to *Bactrocera* had been demonstrated by many previous studies [65–68], which further be verified by our results. Besides, *S. atilius* and *T. femoralis* were closely related and they gathered to form a sister group to *P. utilis*. The evolutionary status and phylogenetic relationship of *S*. *atilius* firstly were investigated and clarified. Although the known knowledge from *S*. *atilius* were still limited, our study would be helpful to provide a molecular basis for the classification and phylogeny of *S*. *atilius* including other Tephritidae species.

## Conclusion

In conclusion, the mitochondrial genome of *S*. *atilius*, which was the first mitochondrial genome of Sphaenisscus, showed high conservation in terms of gene size, organization, AT bias, and secondary structures of tRNAs. The evolutionary status and phylogenetic relationship of *S*. *atilius* were investigated and clarified, revealing that *S. atilius* and *T. femoralis* were closely related and form a sister group to *P. utilis*. These results provided essential data for the phylogenetic and evolutionary analysis of *S*. *atilius*.

## Author Contributions

**Data curation:** Shibao Guo, Nan Song.

**Formal analysis:** Fangmei Zhang, Nan Song.

**Investigation:** Junhua Chen

**Methodology:** Fangmei Zhang, Nan Song.

**Funding acquisition:** Shibao Guo.

**Supervision:** Junhua Chen

**Writing-original draft:** Shibao Guo, Fangmei Zhang.

**Writing-review & editing:** Shibao Guo, Fangmei Zhang, Nan Song.

## Funding

This study was supported by Special funds for Henan Provinces Scientific and Technological Development Guided by the Central Government (Z20221341063), Natural Science Foundation of Henan Province (No. 212300410229), Key Project for University Excellent Young Talents of Henan Province (No. 2020GGJS260), the Project of Science and Technology Innovation Team (No. XNKJTD-007 and KJCXTD-202001). The funders had no role in study design, data collection and analysis, decision to publish, or preparation of the manuscript.

## Competing interests

The authors declare no conflict of interest.

## Notes

### Competing Interest Statement

The authors have declared no competing interest.

